# Functional connectome analyses reveal a highly optimized human olfactory network

**DOI:** 10.1101/843532

**Authors:** T. Campbell Arnold, Yuqi You, Mingzhou Ding, Xi-Nian Zuo, Ivan de Araujo, Wen Li

## Abstract

The olfactory system is uniquely heterogeneous, performing multifaceted functions (beyond basic sensory processing) across diverse, widely distributed neural substrates. While knowledge of human olfaction continues to grow, it remains unclear how the olfactory network is organized to serve this unique set of functions. Leveraging a large and high-quality resting-state functional magnetic resonance imaging (rs-fMRI) dataset of nearly 900 participants from the Human Connectome Project (HCP), we identified a human olfactory network encompassing cortical and subcortical regions across the temporal and frontal lobes. Highlighting its reliability and generalizability, the connectivity matrix of this olfactory network mapped closely onto that extracted from an independent rs-fMRI dataset. Graph theoretical analysis further explicated the organizational principles of the network. The olfactory network exhibits a functionally advantageous modular composition of three (i.e., the *sensory*, *limbic*, and *frontal*) subnetworks and demonstrates strong small-world properties, high in both global integration and local segregation (i.e., circuit specialization). This network organization thus ensures the segregation of local circuits, which are nonetheless integrated via connecting hubs (i.e., amygdala and anterior insula), thereby enabling the specialized, yet integrative, functions of olfaction. In particular, the degree of local segregation positively predicted olfactory discrimination performance in the independent sample. In sum, an olfactory functional network has been identified through the large HCP dataset, affording a representative template of the human olfactory functional neuroanatomy. Importantly, the topological analysis of the olfactory network provides network-level insights into the remarkable functional specialization and spatial segregation of the olfactory system.

**Significance Statement:** Olfaction is an intriguing multifunctional system, playing key roles in regulating emotions, autonomic tone, and feeding, beyond basic sensory perception. However, it is unclear how the neuroanatomy of olfaction is organized in humans to subserve these functions. Functional connectivity analysis of the HCP dataset combined with graph theoretical analysis revealed an optimized large-scale network consisting of three subnetworks—the sensory, limbic, and frontal subnetworks. Distributed across frontal and temporal lobes in well segregated fashion, these olfactory structures are also highly integrated, linked through hub nodes of the amygdala and anterior insula. Our independent dataset replicated the HCP-derived olfactory network and, importantly, highlighted a direct association between the degree of network segregation and olfactory perception.

## Introduction

The olfactory system is uniquely heterogeneous, with functions that extend well beyond basic sensory processing to include domains of emotion, neuroendocrine, and homeostasis (Shipley and Ennis, 1996; Shepherd, 2004). Accordingly, the olfactory neuroanatomy involves widely distributed cortical and subcortical structures, exhibiting a high degree of functional specialization and spatial segregation (Kjelvik et al., 2012; Mainland et al., 2014; Kondoh et al., 2016; Zou et al., 2016). The human olfactory system comprises a set of primary (receiving direct bulbar input) and secondary olfactory regions in the temporal and frontal lobes (Carmichael et al., 1994; Gottfried and Zald, 2005; Zelano and Sobel, 2005; Seubert et al., 2013). Complex, large-scale networks integrated across distributed structures have been increasingly recognized as the fundamental organizational architectures and operational units of the brain (Varela et al., 2001; Fox and Raichle, 2007; Yuste, 2015). Here, we sought a network-level understanding of the human olfactory system—how are the olfactory regions organized to support diverse, yet highly integrated, functions?

Resting-state functional magnetic resonance imaging (rs-fMRI) in humans has revealed robust inter-regional coupling of spontaneous fMRI signal fluctuations underlying intrinsic functional connections (Biswal et al., 1995). This research has identified stable large-scale resting-state networks (RSNs), including networks of the physical (visual, auditory, and somatosensory) senses, but the olfactory network remains elusive. The largely subcortical composition of the olfactory system, with many loci at the air-tissue interface, has presented a serious challenge to olfactory network identification, especially for unguided, whole-brain rs-fMRI connectivity analysis. However, important insights into the olfactory network have been gained by targeting olfactory regions of interest (ROIs) (Plailly et al., 2008; Krusemark and Li, 2012; Krusemark et al., 2013; Sunwoo et al., 2015; Kollndorfer et al., 2015; Novak et al., 2015; Karunanayaka et al., 2017; Milardi et al., 2017; Fjaeldstad et al., 2017; Cecchetto et al., 2019), especially in combination with network-science analysis (Meunier et al., 2014; Royet et al., 2011; Ripp et al., 2018; Zhou et al., 2019). The olfactory ROIs are fairly reliably identified, but inconsistencies in network composition and connections also abound in this literature (Fjaeldstad et al., 2017; Cecchetto et al., 2019).

Disparities in tasks employed in previous studies present a major source of inconsistency by engaging different regions and pathways. In comparison, rs-fMRI connectivity analysis is fairly immune to such confounds and has indeed revealed more reliable and robust connections than task-positive analyses (Sporns, Tononi & Edelman, 2000; Braun et al., 2012; Cao et al., 2014; Zuo et al., 2019). Another major source of inconsistency concerns the idiosyncratic nature of human olfactory perception and neuroanatomy (Richardson and Zucco, 1989; Krusemark and Li, 2012). That is, to produce a reliable and representative depiction of the system, sufficiently large samples are required to overcome individual variability, but the extant studies have been of modest sample sizes. To address these issues, we leveraged an extraordinary rs-fMRI dataset from the Human Connectome Project (HCP), consisting of nearly 900 individuals from diverse ethnicity/race (S900 data release; Van Essen et al., 2013), and combined ROI-based and whole-brain RS connectivity analysis to delineate the human olfactory network. Furthermore, to demonstrate the generalizability and reproducibility of this delineation, we repeated the analysis in an independent sample from our lab.

After defining the olfactory network composition, we attained insights into the functional organization of the olfactory network using meso-scale network analysis (Fortunato, 2010; Meunier et al., 2010; Karrer and Newman, 2011). Graph theory analysis represents a chief model for meso-scale network architecture (Bullmore and Sporns, 2009; Power et al., 2011) and was applied to explicate the organizational principles of the olfactory network. Importantly, we administered an olfactory discrimination task in the independent sample and correlated the performance with graph theory metrics to test the hypothesis that the olfactory network is optimally organized to facilitate olfactory perception.

## Methods

### Main Study (the HCP dataset)

#### Participants

Participants for the main study were obtained from the open access HCP (Human Connectome Project) S900 data release (Van Essen et al., 2013). The full S900 release contains fMRI (functional magnetic resonance imaging) scans for 897 individuals; our analysis included the 812 subjects, who had all four resting-state scans and a voxel selection masks serving to remove low signal-to-noise ratio (SNR) voxels. To ensure adequate and comparable fMRI signal strengths across the ROIs (many of which are located in areas that are highly susceptible to signal dropout and artifact), we further excluded participants who had (1) less than 50 voxels in any ROI (*n* = 2); (2) a majority (> 60%) of voxels in an anatomical ROI missing from the functional scans (*n* = 57); or (3) a SNR of any ROI that was 3 *SD*s below the sample mean (*n* = 60). The olfactory tubercle, a very small structure in humans, was exempted from this exclusion. Based on these criteria, 84 participants were excluded, resulting in a final sample of 728 participants (405 females; age: 28.8 ± 3.7 years).

#### Image Acquisition and Preprocessing

Resting-state scans were collected on a 3T Siemens Skyra MRI scanner and 32-channel head coil using a gradient-echo EPI (Echoplanar Imaging) sequence. The imaging parameters were TR/TE = 720/33 ms; flip angle = 52°; field of view = 208 × 180 mm; matrix size = 104 × 90; slice thickness = 2.0 mm, 72 slices, 2.0 mm isotropic voxels; multiband factor = 8. Two runs (one Right-Left and one Left-Right phase coding) were collected on each of two consecutive days. Each run contained 1200 volumes. Note, due to high signal dropout in APC and OFC regions in the Left-Right phase-encoding runs, only the Right-Left phase-encoding runs were included for these regions. The large number of scans for the Right-Left encoding runs (*n* = 2400) would nonetheless ensure sufficient data for connectivity analysis concerning these regions. A T1-weighted structural image was collected on day one (0.7 mm isotropic voxels).

All images were preprocessed according to the HCP minimal preprocessing pipeline including artifact removal, motion correction, fieldmap correction, high-pass filtering, and normalization to the MNI (Montreal Neurological Institute) template. Full details on the HCP data set and the preprocessing pipeline can be found in the S900 release manual and in previously published overviews of the HCP procedures (Glasser et al., 2013; Smith et al., 2013). Further motion correction procedures in preparation for connectivity analysis are described below.

#### Brain Parcellation

The brain was parcellated based on a version of the Automated Anatomical Labeling (AAL) atlas (Tzourio-Mazoyer et al., 2002) that has been further subdivided into 600 cortical regions of roughly similar size through iterative bisection of larger regions (Hermundstad et al., 2013). Additionally, 28 cerebellum parcels were added (Diedrichsen et al., 2009). Several key olfactory regions, which are subcortical and/or not well defined in the AAL atlas, were drawn on the group-average anatomical T1 (HCP S900 release) in MRIcro (Rorden and Brett, 2000) in reference to a human brain atlas (Mai et al., 2008). These regions include the anterior and posterior piriform cortex (APC and PPC), amygdala (AMY), anterior and posterior hippocampus (aHIP and pHIP), entorhinal cortex (ENT), olfactory tubercle (OTB), nucleus accumbens (NAcc), and hypothalamus (HYP). The olfactory orbitofrontal cortex (OFC) region was defined by a 12 × 12 × 10 mm^3^ box centered around the putative olfactory OFC centroid [25, 35, −14] (Gottfried and Zald, 2005). The eight insula parcels in the 600 parcellation set were merged into four regions, posterior, dorsal, ventral, and anterior insula (INSp, INSd, INSv, and INSa, respectively), to coincide with the human insular functional anatomy as delineated in a neuroimaging meta-analysis (Christopher et al., 2014). Voxels included in both the original AAL parcels and the drawn/adjusted regions were assigned to the latter, generating a final brain parcellation of 627 regions. The atlas was generated using the SPM (http://www.fil.ion.ucl.ac.uk/spm/software/spm12/) software packages for Matlab (Mathworks, Natick, MA) and FreeSurfer (Fischl, 2012).

The mean size of the parcels (excluding large cerebellum parcels) was 266 ± 76 voxels. To address signal dropout and artifacts, we conducted the following voxel removal procedure: (1) A brain mask generated from each participant’s T1 was used to exclude voxels located outside the participant’s brain; and (2) Voxels determined to have a high Coefficient of Variation (COV) were excluded from the parcel (cutoff: COV more than 0.5 standard deviations above the mean within a 5-mm sigma Gaussian neighborhood) (Glasser et al., 2013).

#### Regions of Interest

Twenty-eight regions were targeted as regions of interest (ROIs) based on their possible roles in olfaction: (1) key olfactory areas—five regions receiving direct olfactory bulbar input, including APC, PPC, AMY, ENT, and OTB (Carmichael et al., 1994; Haberly, 2001; Gottfried and Zald, 2005; Seubert et al., 2013) as well as the olfactory orbitofrontal cortex (Oolf), a well-established region in the human olfactory neuroanatomy (Gottfried, 2010). The anterior olfactory nucleus was not included due to limited accessibility by fMRI. (2) secondary, associative olfactory regions—additional regions in the OFC, insula (INSp, INSd, INSv, and INSa), thalamus (dorsal anterior/THLda; ventral anterior/THLva; dorsal posterior/THLdp; and ventral posterior/THLvp), aHIP, pHIP, NAcc, and HYP (Carmichael et al., 1994; Haberly, 2001; Gottfried and Zald, 2005; Seubert et al., 2013). To have a comprehensive examination of the OFC participation in olfaction, we included the entire OFC (barring the most lateral parts), with a total of 11 parcels. The ventral medial prefrontal cortex was not considered here. Due to signal susceptibility at the tissue-air conjunction, we observed considerable signal dropout in the left hemisphere at the bottom of the frontal lobe and the fronto-temporal junction, affecting key regions of interest (in the posterior OFC and APC). In light of predominantly ipsilateral olfactory pathways (Shipley and Ennis, 1996) and similar functional connectivity between the hemispheres (Zhou et al., 2019), we confined our olfactory network analysis to the right hemisphere where fMRI signals at the susceptible areas were well preserved. **Table 1** shows centroid coordinates, sizes, and mean correlation coefficients (among the ROIs and among the whole brain) for all ROIs. **Figure 1** illustrates the procedure **(A)** and anatomical locations of all ROIs before and after voxel removal **(B)**.

**Figure 1.**
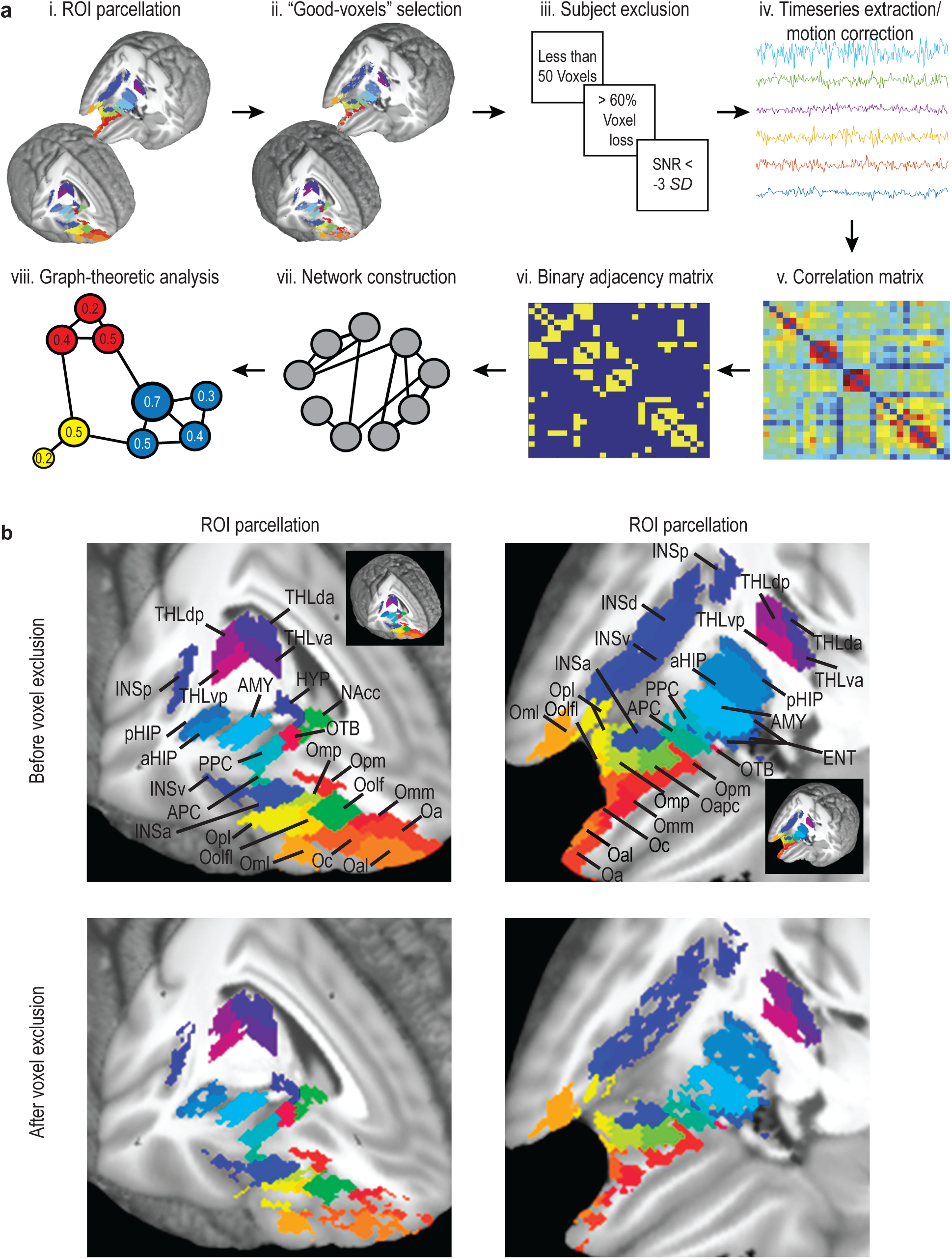
Procedures. **(A)** Schematic illustration of analysis pipeline. (i) A total of 28 ROIs were defined. (ii) An automated procedure based on Coefficient of Variation (COV) removed voxels contaminated with artifacts from the ROIs. (iii) Participant exclusion based on three exclusion criteria. (iv) Timeseries data extraction from the ROIs. (v) A 28-by-28 correlation matrix compiled based on pair-wise correlations across the ROIs. (vi) A binary adjacency matrix constructed with supra- and sub-threshold connections. (vii) Suprathreshold connections chosen to form the olfactory network. (viii) Graph-theoretical analyses performed to characterize the organization of this network. (**B)** 3D display of ROIs before (top row) and after (bottom row) voxel removal. Insets illustrate the underlying ROIs in 3D whole brain images with parts of dorsolateral frontal and temporal lobes removed.

**Table 1.** ROI centroid coordinates, sizes, and mean (whole-brain) correlation coefficients

| **ROI** | **x** | **y** | **z** | **voxels** | ***r*** |
| --- | --- | --- | --- | --- | --- |
| Oolf | 25 | 35 | -14 | 180 | 0.25 |
| APC | 24 | 10 | -20 | 143 | 0.22 |
| PPC | 24 | 4 | -17 | 137 | 0.22 |
| AMY | 21 | -5 | -20 | 377 | 0.30 |
| aHIP | 26 | -14 | -21 | 312 | 0.28 |
| pHIP | 28 | -28 | -9 | 311 | 0.28 |
| ENT | 18 | -4 | -29 | 309 | 0.25 |
| INSa | 35 | 21 | -6 | 449 | 0.30 |
| INSd | 39 | 8 | 6 | 421 | 0.27 |
| INSv | 43 | 2 | -5 | 513 | 0.29 |
| INSp | 37 | -15 | 12 | 368 | 0.29 |
| HYP | 5 | -3 | -12 | 107 | 0.22 |
| THLda | 11 | -14 | 13 | 263 | 0.25 |
| THLva | 10 | -13 | 2 | 276 | 0.27 |
| THLdp | 16 | -24 | 11 | 238 | 0.24 |
| THLvp | 14 | -25 | 2 | 280 | 0.27 |
| OTB | 14 | 4 | -14 | 43 | 0.13 |
| Omm | 16 | 41 | -21 | 204 | 0.24 |
| Opm | 17 | 25 | -21 | 222 | 0.27 |
| Oa | 15 | 56 | -17 | 250 | 0.23 |
| Oc | 28 | 43 | -17 | 219 | 0.23 |
| Oal | 24 | 57 | -14 | 252 | 0.20 |
| Oml | 42 | 38 | -12 | 250 | 0.26 |
| Oolfl | 33 | 34 | -15 | 183 | 0.24 |
| Opl | 39 | 26 | -15 | 233 | 0.25 |
| Omp | 31 | 24 | -21 | 160 | 0.22 |
| Oapc | 23 | 18 | -23 | 134 | 0.22 |
| NAcc | 10 | 11 | -8 | 116 | 0.22 |

#### Timeseries extraction and artifact removal

BOLD values for each scan were averaged across all voxels within an ROI for each resting-state run, resulting in four sets of 627 ROI timeseries each consisting of 1200 scans per participant. To further remove artifacts that could contribute to spurious RS activity variance (Fox et al., 2009; Power et al., 2012), several additional preprocessing steps were implemented. These steps included (1) mean centering and whitening of timeseries before concatenating runs of the same phase-encoding; (2) applying a temporal bandpass (.01-.08 Hz) filter (Biswal et al., 1995); (3) running a general linear model (GLM) to regress out head motion (Satterthwaite et al., 2013; Yan et al., 2013). The model contained 24 nuisance variables (Friston et al., 1996), including six head motion parameters from the current time point, the six parameters from the previous time point, and the squared values of the first twelve parameters; (4) and scrubbing of “spikes” containing significant motion based on framewise displacement index (FDi) defined as [FDi= |∆dix| + |∆diy| + |∆diz| + |∆αi| + |∆βi| + |∆γi|]; Scans with FDi over 0.5 mm were classified as spikes in movement and removed (Power et al., 2012; Spielberg et al., 2015).

#### Network Construction

Based on the extracted timeseries, Pearson’s correlation coefficients were calculated for each ROI pair and used to construct a 28 × 28 correlation matrix for each participant. These correlation coefficients were then Fisher Z-transformed, and for each pair, the coefficient was submitted to a *t*-test against its global baseline connectivity. This global baseline connectivity was defined for each pair as the average correlation coefficient (Fisher-transformed) of the two regions with all other (625) regions in the whole brain. A correlation in the top 5% of the 625 connections for each node was considered as significant (Watrous et al., 2013; Schedlbauer et al., 2014). A binary adjacency matrix, denoting supra- and sub-threshold connections, was thus constructed. These procedures are illustrated in **Figure. 1A** (v-vi).

For network construction, we first identified suprathreshold connections among the six key olfactory areas (APC, PPC, AMY, ENT, OTB, and Oolf,). Secondary regions (i.e., regions receiving direct input from the primary regions) were defined as any of the remaining 22 ROIs that had at least one connection to one of the key olfactory regions and were thus admitted into the olfactory network.

#### Graph-theoretic Analysis

To characterize the olfactory network, graph-theory-based analysis was performed on the olfactory network constructed above using the Brain Connectivity Toolbox (Rubinov and Sporns, 2010).

##### Modularity

Modularity maximization algorithms seek to find a division of nodes into subnetworks, or modules, which maximizes the number of intra-module connections while minimizing the number of inter-module connections (Newman, 2006). We performed modularity analysis using the Louvain algorithm (Blondel et al., 2008) as well as the Girvan-Newman--algorithm (Girvan and Newman, 2002) with 10,000 iterations. Modularity index (*Q*) ranges [−1/2,1], with a cutoff score of 0.3 to indicate strong existence of subnetworks(Girvan and Newman, 2002). The reliability of modularity was further tested against randomly reassigned connections (*Q*_*rand*_) based on 10,000 permutations. This distribution was used to calculated the Z-scored modularity, with a score greater than 3 indicating the existence of subnetworks (Fortunato, 2010; Kinnison et al., 2012). The network topology was then illustrated using the Gephi software with the Force Atlas and the expansion tool to increase spacing (Bastian et al., 2009; Zuo et al., 2012).

##### Metrics of network functionality

Several other graph theory metrics, concerning global and local qualities, were extracted to characterize the olfactory network. Small-world network organization is deemed to be highly efficient for spreading information and conserved across species (Rubinov and Sporns, 2010; Bota et al., 2015; Betzel and Bassett, 2018). We thus assessed small-world properties of the olfactory network by extracting two key markers—the global efficiency (*G*, indexing network integration and global communication) (Latora and Marchiori, 2001) and clustering coefficient (*C*, indexing segregation or presence of local clusters) (Watts and Strogatz, 1998). These indices were further contrasted with the average for 10,000 random reassignments of the edges in the network (*G*_*rand*_ and *C*_*rand*_). In a small-world network, global efficiency should be close to that of a random network characterized by a multitude of (random) connections whereas the clustering coefficient should be higher than that of a random network (as random connections are less likely to form clusters).

We also identified local regions key to the organization of the olfactory network. First, we assessed each region’s critical contribution to the network using targeted node deletion. We iteratively removed each of the regions (nodes) and examined the percentage reduction in global efficiency of the network (Bassett and Bullmore, 2006; van den Heuvel and Sporns, 2013). Second, we evaluated each region’s centrality (aka, “hubness”) in the network using three key metrics of centrality—node degree, betweenness centrality, and closeness centrality (Bota et al., 2015). These measures were further aggregated to generate an overall ranking score and a composite score by averaging across the z-scored values (Sporns et al., 2007). Node degree refers to the number of direct connections between a node and any other nodes in the network. Betweenness centrality represents the number of shortest paths that travel through a given node (Freeman, 1978), calculated as the fraction of all shortest paths in the network that contain a given node. Closeness centrality is a measure of the distance from a node to other nodes, calculated as the reciprocal of the average path length between a node and all other nodes in the network (Freeman, 1978). Lastly, we determined whether a hub was a connector or provincial hub using the participation coefficient (*P*), which indexes the diversity of module connections of a given node (Guimerà and Amaral, 2005).

#### Control Analyses: *(dis)Connectivity of the olfactory network with the occipital visual cortex*

To ascertain the validity and specificity of the olfactory network, we constructed a binary connectivity matrix between the olfactory network regions and occipital visual cortices. The occipital lobe was parcellated into a total of 28 parcels located in the Calcarine (8 parcels), Cuneus (4 parcels), superior occipital gyrus (4 parcels), middle occipital gyrus (8 parcels), and inferior occipital gyrus (4 parcels) as defined in the 600 region parcellation of the AAL atlas (Tzourio-Mazoyer et al., 2002). These regions were parcellated into approximately equal sizes by the same method described in the Brain Parcellation section above.

### Validation and Extension Study (the Independent Dataset)

As a validation of the olfactory network defined by the large HCP dataset, we applied this olfactory network to an independent resting-state fMRI dataset collected in our lab. To further link the olfactory network functionality to olfactory performance, we administered an olfactory discrimination task immediately after the resting-state fMRI scan and correlated participants’ global small-world metrics of their olfactory network with their task performance.

#### Participants

Thirty-three healthy participants took part in the study in exchange for course credit or monetary compensation. All participants were right-handed with normal olfaction, which was determined based on participants’ self-reported sense of smell and objective assessment (including odor intensity and pleasantness ratings) during a lab visit. Individuals showing aberrant olfactory performance or with nasal infections/allergies were excluded from participating in the study. Participants were also screened for any history of severe head injury, psychiatric or neurological disorders or current use of psychotropic medication. All participants provided informed consent to participate in the study, which was approved by the University of Wisconsin-Madison Institutional Review Board. One participant was excluded due to metal artifact, leaving 32 participants (13 males; age 19.9 ± 2.0 years, range 18–30) in the final sample.

#### Odor Discrimination Task

Following a resting-state fMRI scan (detailed below), we administered a 2-alternative-forced-choice (2AFC) odor discrimination task, where five binary odor mixtures with systematically varying proportions of acetophenone (“almond”, 5% l/l diluted in mineral oil) and eugenol (“clove”, 18% l/l) were presented. The five odor mixtures contained acetophenone/eugenol ratios of 80/20%, 60/40%, 50/50%, 40/60%, and 20/80%, respectively. Concentrations for the two odorants were determined through careful piloting to ensure equivalent perceived intensity. Prior to the test, participants were presented with acetophenone and eugenol in their original concentrations, which were labeled as Odor A and Odor B (the order was counterbalanced across participants). During the task, participants sniffed an odor mixture and indicated “Odor A” or “Odor B” using a button press. There were 15 trials for each mixture, randomly intermixed with a stimulus onset asynchrony (SOA) of 14.1 ms.

Odor mixtures were delivered at room temperature using a sixteen-channel computer-controlled olfactometer (airflow set at 1.5L/min). When no odor was being presented, a control air flow was on at the same flow rate and temperature. This design permits rapid odor delivery in the absence of tactile, thermal, or auditory confounds (Lorig et al., 1999; Krusemark et al., 2013; Novak et al., 2015). Stimulus presentation and response recording were executed using Cogent software (Wellcome Department of Imaging Neuroscience, London, UK) as implemented in Matlab.

#### Resting-state MRI

##### Image Acquisition

All participants underwent a 10-minute rs-fMRI scan (with eyes open and fixated on the central crosshair) before the odor discrimination task. Gradient-echo T2-weighted echoplanar images (255 scans) were acquired with blood-oxygen-level-dependent (BOLD) contrast on a 3T GE MR750 MRI scanner, using an eight-channel head coil with sagittal acquisition. Imaging parameters were TR/TE: 2350/20 ms; flip angle: 60°, field of view 240 mm, slice thickness 2 mm, gap 1 mm; in-plane resolution/voxel size 1.72 × 1.72 mm; matrix size 128 × 128. A high-resolution (1 × 1 × 1 mm^3^) T1-weighted anatomical scan was also acquired. Lastly, a field map was acquired with a gradient echo sequence.

##### Image Analysis

Imaging data (after removal of the first 6 dummy scans) were preprocessed in SPM12 (http://www.fil.ion.ucl.ac.uk/spm/software/spm12), including slice-time correction, spatial realignment, field-map correction, and normalization to MNI template (2 × 2 × 2 mm voxels) using Diffeomorphic Anatomical Registration Through Exponentiated Lie algebra (DARTEL) (Ashburner, 2007). Based on the olfactory network identified through the HCP data, we focused on the 22 ROIs in the olfactory network, which were defined in the main study. We then applied the same artifact and voxel removal steps as described in the **Timeseries extraction and artifact removal** section above.

#### Correlation Matrix and Graph-theoretic Metrics (Weighted)

To examine individual-level graph metrics and associate them to individual differences in olfactory discrimination, we applied weighted graph theoretic analysis. Specifically, we calculated correlation coefficients for timeseries of any two regions, generating a correlation matrix (22 × 22) for each individual participant. We used the absolute value of correlations to reflect connectivity strength. The matrix was then multiplied with the binary matrix defined by the HCP dataset, constructing a sparse weighted matrix for each participant. The global graph metrics related to small-world-ness, the global efficiency and the clustering coefficient, were calculated based on this sparse weighted matrix (using absolute value weights).

#### Correlational Analysis

Given the likely non-Gaussian distribution of correlation matrices, we computed the non-parametric Spearman correlation coefficient (Rho), and the statistical significance threshold was determined using null-model permutation tests (*n* = 10,000; *p* < .05 was set at 95^th^ percentile). To assess reliability and generalizability of the HCP-based olfactory network, the group correlation matrix of the independent sample was correlated with the group correlation matrix of the HCP sample. We further examined the relationship between olfactory network organization and olfactory perceptual performance. To index basic olfactory discrimination, we applied signal detection theory analysis on the 2AFC performance and extracted *d’* (Z_hit_− Z_false alarm_) based on their responses to the dominantly acetophenone (80/20%) and dominantly eugenol (20/80%) mixtures. Each participant’s *d*’ score and their graph metrics (global efficiency and clustering coefficient) were then entered into Spearman correlational analysis.

## Results

### The olfactory functional network

According to the network construction procedure described above, we identified the olfactory network consisting of the six key olfactory ROIs and 16 additional parcels showing supra-threshold connectivity with one of the key regions (Fig. 2**A** & **B**). These 16 regions included the nucleus accumbence (NAcc), hypothalamus (HYP), both anterior and posterior parcels of the hippocampus (aHIP and pHIP), all four parcels of the insular cortex (INSa, INSp, INSv, and INSd), the ventral posterior thalamus (THLvp), and seven additional OFC parcels (located in the posterior and middle OFC). As illustrated in the weighted binary connectivity matrix (Fig. 2A), these 22 regions of the network were closely connected, constituting 28.6% (66/231) of all possible pair-wise connections. By contrast, exhibiting high modality-specificity, this olfactory network showed no suprathreshold inter-modality connections with occipital visual areas at 5% and only sparse ones at connection densities set at 10% and 15% (**Fig. 2A)**.

**Figure 2.**
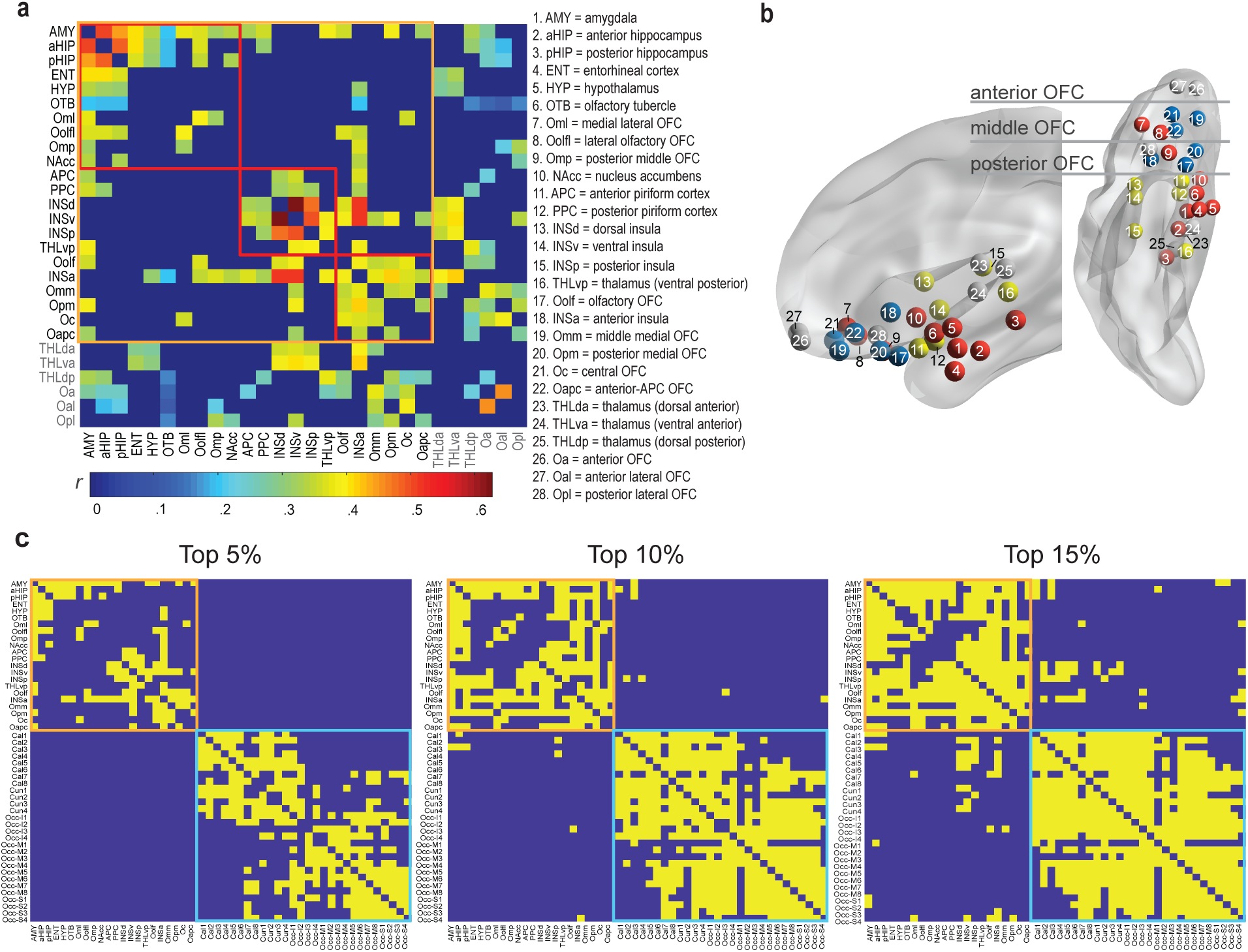
The olfactory network. **(A)** A weighted sparse 28-by-28 correlation matrix of group average Pearson’s *r’*s for all suprathreshold pairs. ROIs included in the olfactory network are enclosed in the orange box, with the three identified modules (subnetworks) enclosed in the red boxes. The table lists the region names in correspondence to the ROI/node numbers. **(B)** A transparent brain model (in sagittal and axial views) with ROIs (nodes) for the three modules coded in three respective colors. Grey nodes are ROIs not accepted into the olfactory network. **(C)** A binary connectivity matrix reveals suprathreshold connections (shown in yellow) across the olfactory network nodes (22 parcels, enclosed in the orange box) and occipital visual cortical regions (28 parcels, enclosed in the cyan box) at three cutoff levels (top 5, 10, and 15%). The visual regions (Cal = Calcarine gyrus, Cun = Cuneus gyrus, Occ-I = Occipital inferior gyrus, Occ-M = Occipital middle gyrus, Occ-S Occipital superior gyrus) were strongly interconnected and relatively disconnected from the olfactory nodes.

### Network organization and characteristics

Graph-theoretic modularity analysis revealed a strong modular organization of the olfactory network, with a reliable composition of 3 modules/subnetworks (*Q*=.30, *Z*=4.99, p<0.001; **Fig. 3A**). As illustrated in Figure 3A, the three modules/subnetworks could be characterized as *i.* the “olfactory sensory subnetwork” consisting of APC, PPC, and three insular parcels (including the ventral, dorsal and posterior, but not the anterior, parcels), in addition to the ventral posterior thalamus on the periphery; *ii.* the “olfactory limbic/paralimbic subnetwork” consisting of the amygdala, olfactory tubercle, and hippocampus at the center, in addition to the entorhinal cortex, hypothalamus, nucleus accumbens, and three OFC parcels on the periphery; and *iii.* the “olfactory frontal subnetwork” consisting of the anterior insula and olfactory OFC at the center and four additional OFC parcels on the periphery.

**Figure 3.**
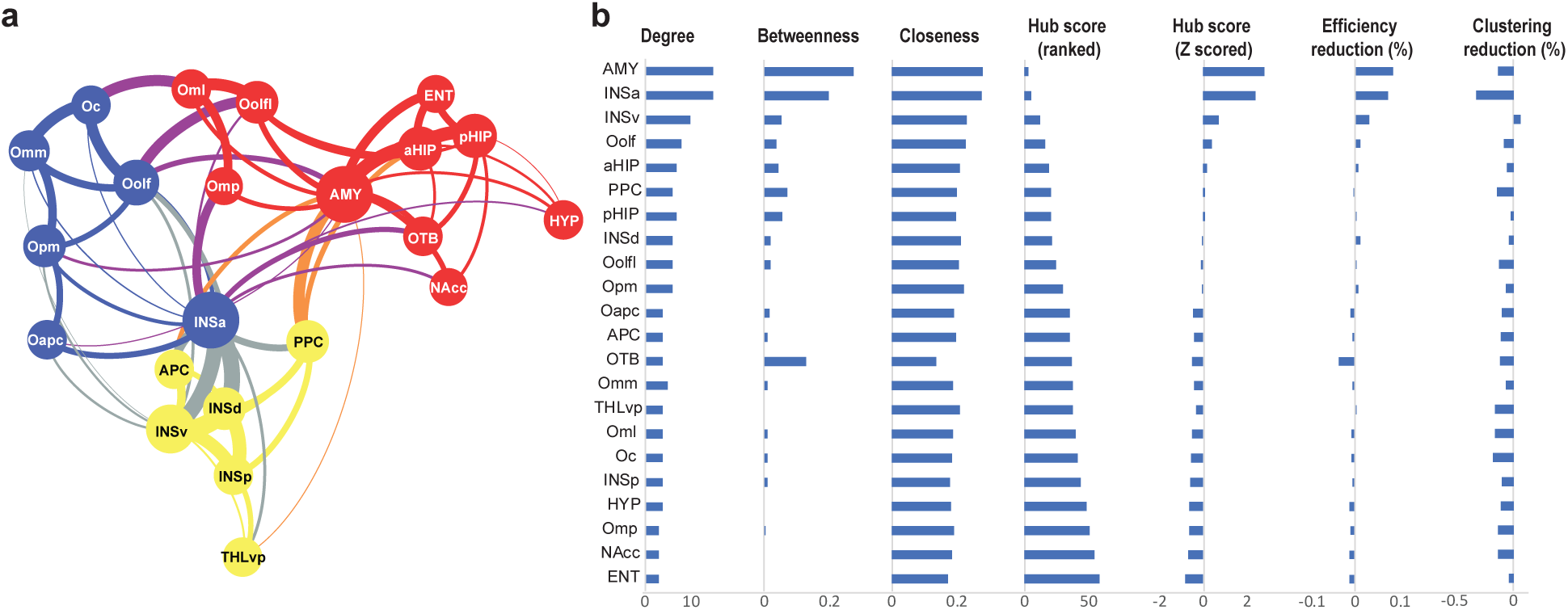
Local network metrics. **(A)** Topology of the olfactory network. The three modules/subnetworks are indicated by the three colors of the circles. Line thickness indicates connection strength (mean correlation coefficients), and node size reflects connection density (number of connections). **(B)** Hubness of a node as reflected by composite hub ranking and composite hub Z-scores. The three centrality indices— node degree, betweenness and closeness centrality)—are also displayed. The AMY and INSa separated from the other nodes as major hubs of the network. Changes in global efficiency of the olfactory network following a node removal were small except for the AMY and INSa nodes, which resulted in 8.5% and 7.3% reductions in global efficiency, respectively.

These subnetworks were connected via multiple between-module connections. Based on a set of graph theoretic metrics of node centrality (i.e., degree, betweenness and closeness), the amygdala and anterior insula stood out as two major hubs of the network (Fig. 3B), whose participant coefficient values (.56 and .67, respectively) also indicated that they were connecting hubs between subnetworks (Fig. 3A). Graph-theoretic analysis further assessed small-world-ness of the olfactory network based on two defining features: Global Efficiency (*G*, indicating network integration and global communication) and Clustering (*C*, indicating segregation of local clusters). Relative to random networks (based on 10,000 random permutations of the olfactory network connections) characterized by high efficiency but low clustering values, the olfactory network exhibited a high degree of small-world organization with high global efficiency (*G* = .231, highly comparable to *G*_rand_ = .236) and high clustering (*C* = .186 exceeding *C*_rand_ = .164).

Finally, given the susceptibility of many olfactory regions to neurodegenerative pathological invasion, we assayed the resilience of the olfactory network to local attacks using iterative node removal. The olfactory network sustained minimal impact by the removal of any one node (global efficiency change ranged −3.8% to 3.1%), with the exception of the hub regions (amygdala and anterior insula) whose removal led to modest reductions (8.5% and 7.3%, respectively; Fig. 3B).

### Linking network organization to olfactory perception

To ascertain the functional relevance of the olfactory network organization, we then associated the olfactory network metrics with olfactory performance (i.e., odor discrimination) in an independent sample collected in the lab. First, we validated the olfactory network in this sample: there was strong concordance between the connectivity matrices derived from the HCP and independent samples, Spearman *rho* = .41, *p* < .001 (**Fig. 4A**), in support of the reliability and generalizability of the olfactory network. Next, applying the olfactory network topology defined by the HCP dataset, we extracted each participant’s weighted small-world-ness metrics for the network—weighted global efficiency (*G*_w_) and clustering (*C*_w_). Spearman correlation analyses indicated that the odor discrimination performance, *d*’, correlated significantly with the clustering coefficient (*rho* = .32, *p* < .05; **Fig. 3B**) but not global efficiency (*rho* = .13, *p* =.224).

**Figure 4.**
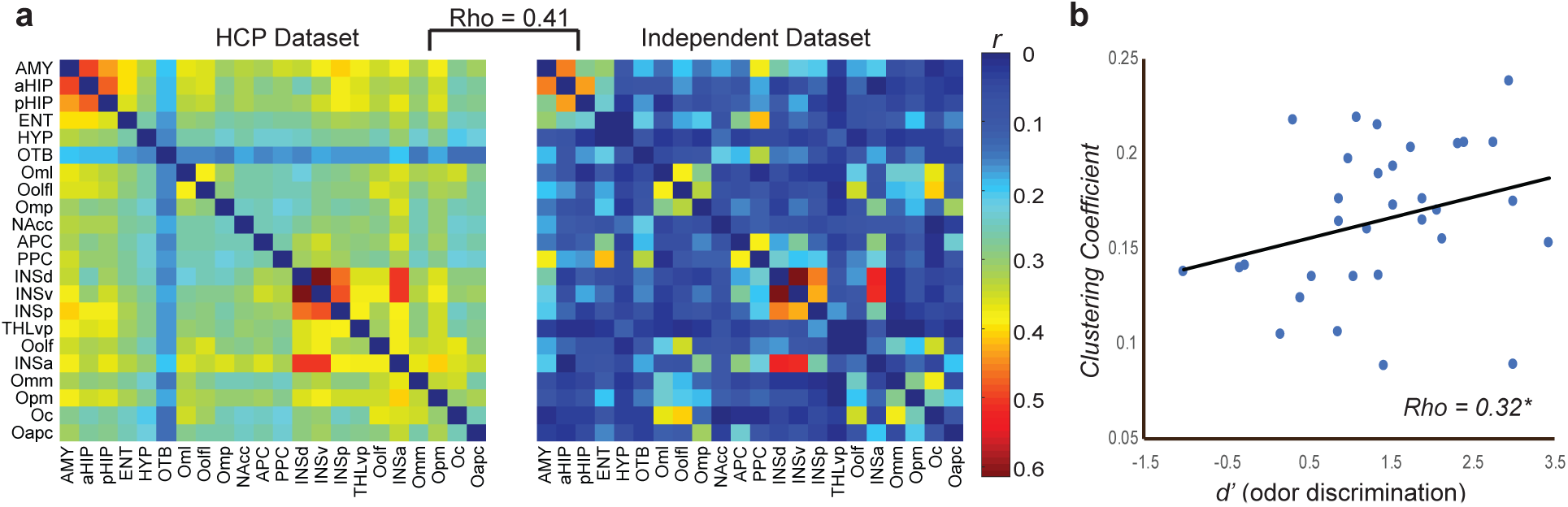
Validation and function of the olfactory network. **(A)** Weighted connectivity matrices of the olfactory network based on the HCP dataset and the independent dataset greatly overlapped; Spearman *rho* = .41, *p* < .001. **(B)** The global network metric of clustering coefficient was positively correlated with olfactory discrimination performance (*d*’), *rho* = .32, *p* < .05.

## Discussion

Combining ROI and whole-brain analyses on the S900 HCP rs-fMRI dataset, we identified a human olfactory functional network of 22 interconnected parcels. Akin to the extraordinary size of the dataset that is conducive to high generalizability, this network demonstrated a strong concordance with the one extracted from our independent dataset. Graph theoretical analysis of the olfactory network further revealed an advantageous modular composition of three subnetworks—the sensory, limbic, and frontal subnetworks. The olfactory network also exhibited strong small-world properties, high in both global integration and local segregation. Importantly, as we hypothesized, the level of local segregation directly predicted odor discrimination performance in the independent sample. In sum, the current study provided a representative description of the human olfactory network and potentially, a template for the functional neuroanatomy of human olfaction. Furthermore, the network topology indicates an optimally organized architecture well suited for the diverse, specialized functions of olfaction.

This olfactory network comprised all ROIs (except for two anterior and one lateral OFC parcels and three thalamus parcels) implicated in rodent and non-human primate neuroanatomy, lending credence to the strong phylogenetic conservation of the olfactory system (Zelano and Sobel, 2005; Gottfried, 2010; Royet et al., 2011; Seubert et al., 2013). Connectivity density of the olfactory network was rather low (28.6%), which is consistent with other sensory networks (Young, 1993; Bassett and Bullmore, 2006) and accords with the evolutionary pressure to keep wiring to minimum to reduce communication cost (Cowey, 1979; Mitchison, 1991). Nonetheless, this connectivity density was in strong contrast with the highly sparse inter-network connectivity between the olfactory network and occipital visual cortices. That is, no inter-network connectivity survived the statistical threshold (top 5%), and only at a lenient threshold (top 10%) did we observe connections between certain visual cortical parcels and the limbic regions of the olfactory network (e.g., amygdala and hippocampus), which are known to receive visual input (Augustine, 1996; Freese and Amaral, 2009). These findings confirm that the olfactory network was modality-specific and indicate a clear preference in connectivity between the primary olfactory structures and the multimodal limbic regions, underscoring their integral participation in the olfactory network.

The role of the thalamus in olfaction has been unclear in the literature. Here, the human olfactory network included only one parcel (in the ventral posterior portion) of the thalamus, which joined the network as a peripheral node. Furthermore, this parcel was connected only with the amygdala and insula, known to receive strong thalamic projections largely conveying sensory input from other modalities (Augustine, 1996; Freese and Amaral, 2009; Gogolla, 2017). These results thus concur with the view of a minor contribution of the thalamus to the olfactory system (Smythies, 1997; Shepherd, 2005) and reinforce the notions of sparse thalamic connections with olfactory cortices and the lack of an obligatory thalamic relay (Price and Slotnick, 1983; Price, 1985). That said, while no corticothalamic connections reached the conventional threshold applied here (top 5%), they emerged at a lenient threshold (top 10%), including connections with the APC and multiple OFC parcels (Fig. 2C). Therefore, these findings permit the possibility that the weak corticothalamic pathways can be strengthened to become functionally relevant with certain task demands. For example, the olfactory-cortex-thalamus-OFC pathway was found to become significant during active attention, thereby engaging the OFC to subserve high-level olfactory processing (Plailly et al., 2008).

As the olfactory system supports not only olfactory sensory perception but also multiple non-sensory functions such as emotion and homeostasis (e.g., neuroendocrine regulation, reproductive response, feeding; Shipley, 1974), the composition of widely distributed cortical and subcortical structures in the olfactory network is consistent with the heterogeneous functions it serves. Accordingly, such diverse functions carried out across a widely distributed network would also demand a highly optimized network organization. Indeed, meso-scale graph theoretical analysis of the network topology confirmed the efficient organization of the olfactory network to suit its remarkable functionality.

First, we observed that the olfactory network had a strong degree of modularity, consisting of three modules with dense intra-module and loose inter-module connections. Modularity is a hallmark feature of optimized networks as formations of self-contained subdivisions effectively reduce overall network complexity and insulate local errors from global network functioning (Ash and Newth, 2007). Not only was the olfactory network compartmentalized into three subdivisions, but also the subdivisions were highly aligned with their diverse yet specialized functions. That is, a sensory subnetwork (comprising APC, PPC, and insula and thalamus parcels) would subserve basic olfactory sensory processing, a limbic subnetwork (comprising limbic ROIs and three lateral OFC parcels) would support emotion and homeostasis, and a frontal subnetwork (comprising OFC parcels and anterior insula) would underpin higher-level, integrative processes.

Second, the olfactory network had a “small-world” quality, a defining feature of a highly optimized network. Specifically, the olfactory network possessed a combination of high global efficiency and high local segregation. As such, the olfactory network assumed efficient communication across the subnetworks to allow for integrative processing while maintaining sufficient segregation to preserve specialized analysis. Evidently, this balance of global integration and subnetwork segregation is well suited for the olfactory system to sustain its diverse but specialized functions in a highly integrated manner.

Third, the olfactory subnetworks were integrated via two key hub structures, the amygdala and anterior insula, akin to their strong connections with temporal and frontal structures (Freese and Amaral, 2009). In fact, given their connections with an extensive web of brain regions, the amygdala and insula have been recognized as central hubs of large-scale neural systems (van den Heuvel and Sporns, 2013; Bickart et al., 2014; Gogolla, 2017). As such, with the intimate participation of the amygdala and insula, the olfactory network is well positioned to summon a high level of global integration. Functionally, the amygdala can relay emotional and homeostatic signals and the insula interoceptive signals to the sensory subnetwork to imbue olfactory perception with rich emotional and homeostatic information. Dovetailing with this organization, alliesthesia, a sensory experience that closely depends on the internal physiological milieu, prevails in olfaction (Cabanac, 1971, 2004; Krusemark et al., 2013). For instance, depending on the level of metabolic energy reserve (hungry or satiated), a food odor, while maintaining its odor identity (as processed in the sensory subnetwork), would take on distinct biological and emotional qualities (appetizing/pleasant or unappetizing/unpleasant) by integrating emotional and physiological signals relayed by the amygdala and insula.

Lastly, the olfactory network would be resilient from local attacks. Complex networks are known to be tolerant to random errors such that a local malfunction would not cause global network dysfunctions (Callaway et al., 2000; Crucitti et al., 2004). Likewise, as we observed here, the removal of a region (other than the hubs) from the olfactory network resulted in minimal loss (< 5%) in global efficiency. Remarkably, to the extent that specific hub failures often result in substantial global deficiency in a network (Callaway et al., 2000; Crucitti et al., 2004), the olfactory network sustained only modest loss (< 10%) in global efficiency (van den Heuvel and Sporns, 2011) with the removal of the amygdala or anterior insula. This level of resilience could be especially valuable for maintaining the overall integrity of the olfactory network as many of its structures, including the hubs (amygdala and insula), are susceptible to pathological invasions by disorders such as Alzheimer’s disease (Herzog and Kemper, 1980; Braak and Braak, 1991; Mesulam, 2015). This resilience can thus explain the fact that early-stage or preclinical patients maintain largely preserved global olfactory functions despite various (and often discrete) perceptual impairments (Doty et al., 1987; Royet, 2001; Djordjevic et al., 2008; Li et al., 2010; Wilson et al., 2010).

The functional relevance of the olfactory network organization was evinced by our independent dataset, where we correlated small-world indices with olfactory discrimination performance. We observed that local segregation, but not global efficiency, was critical for accurate olfactory discrimination. High segregation improves local efficiency and promotes specialized processing within local circuits (Rubinov and Sporns, 2010). Given its heterogeneous composition and the wealth of non-sensory input it receives, it stands to reason that the olfactory network needs to impose a certain level of functional insulation to its sensory subnetwork, thereby ensuring sensory fidelity in basic olfactory perception. That is, odor quality processing in the sensory subnetwork can be insulated against non-sensory influences from the other subnetworks such that olfactory perceptual validity is preserved. Alternatively, leakage from the limbic and frontal subnetworks, especially in individuals with low local segregation, would infuse an odor with hedonic hues or cognitive biases, which dominate and even alter olfactory perception. By this extension, variability in local segregation of the olfactory network could represent a viable network account for the idiosyncrasy of human olfactory experiences.

In conclusion, the human olfactory neuroanatomy involves an evolutionarily conserved, topologically organized large-scale network. The compartmentalization of subnetworks allows for the specialized and yet multifaceted functions of the olfactory system while strong global integration welds the subnetworks to perform integrated processes. Arising from this highly optimized network, are the complex, varied, and almost infinite smells that define human olfaction (Yeshurun and Sobel, 2010; McGann, 2017).

## Acknowledgments

This research was supported by the National Institute of Mental Health R01MH093413 (W.L.).

